# Adversarial generation of gene expression data

**DOI:** 10.1101/836254

**Authors:** Ramon Viñas, Helena Andrés-Terré, Pietro Liò, Kevin Bryson

## Abstract

The problem of reverse engineering gene regulatory networks from high-throughput expression data is one of the biggest challenges in bioinformatics. In order to benchmark network inference algorithms, simulators of well-characterized expression datasets are often required. However, existing simulators have been criticized because they fail to emulate key properties of gene expression data.

In this study we address two problems. First, we propose mechanisms to faithfully assess the realism of a synthetic gene expression dataset. Second, we design an adversarial simulator of expression data, gGAN, based on a Generative Adversarial Network. We show that our model outperforms existing simulators by a large margin, achieving realism scores that are up to 17 times higher than those of GeneNetWeaver and SynTReN. More importantly, our results show that gGAN is, to our best knowledge, the first simulator that passes the Turing test for gene expression data proposed by Maier et al. (2013).

## 1 Introduction

Over the last two and a half decades, the emergence of technologies such as spotted microarrays (Schena et al., 1995) and RNA-seq (Mortazavi et al., 2008) has enabled to accurately measure the expression levels of thousands of genes. In parallel, the interest of the research community for analyzing these profiles has increased over years, resulting in the development of several bioinformatics tools to find recurring patterns in expression data. Concretely, an important effort has been put into reverse engineering gene regulatory networks (GRNs) for a better understanding of the complex interactions that occur among genes (Yu et al., 2004; Margolin et al., 2006; Yip et al., 2010; among others).

However, relatively little effort has been devoted to benchmarking the performance of bioinformatics methods that infer the structure of GRNs, along with their dynamical properties, from high-throughput experimental data. Evaluating these algorithms is usually a challenging task, because we often lack well-understood biological networks to use as gold standards. When information about the gene regulatory interactions is not available, assessing the performance of network inference algorithms requires repeatedly testing them on large, high-quality datasets derived from well-characterized networks (Van den Bulcke et al., 2006). Unfortunately, datasets of an appropriate size are usually unavailable. Under this scenario, there is a clear need for simulators of high-quality expression datasets.

Generating realistic expression data is a challenging task for two reasons. First, the number of possible GRNs grows exponentially with the number of genes. This poses a major challenge to model the complex regulatory interactions that occur among genes, as these networks are usually unknown. Second, it is difficult to determine to which extent the expression data generated by a simulator is realistic. Unlike in other domains such as image generation, wherein one can empirically assess whether an image is realistic, evaluating the realism of a gene expression dataset is complex because we often lack reliable gold standards and we do not have an intuitive understanding of high-dimensional expression data.

This work aims to address these two challenges. First, we develop our own novel quality assessment measures to evaluate the quality of synthetic datasets. In particular, we check whether cohorts of synthetically generated expression data match several relevant statistical properties of the real data. Second, we implement an adversarial simulator based on an extension of generative adversarial networks (GAN, Goodfellow et al., 2014; WGAN-GP, Gulrajani et al., 2017). This framework describes a method for estimating a generative model by playing a two-player game, wherein the first player learns to generate samples from a particular distribution, while the second tries to discriminate them from samples coming from the true data distribution. This novel deep learning framework has shown promising results for tasks such as image generation (Karras et al., 2017) and, to our knowledge, GANs have not yet been applied successfully to build a simulator of gene expression data.

## 2 Methods

### 2.1 Evaluating artificial gene expression data

Assessing to which extent simulators are able to generate realistic datasets is a difficult task, since we often lack reliable gold standards and we do not have an intuitive understanding of high-dimensional expression data. Maier et al. (2013) propose a way of characterizing several statistical properties of gene expression datasets. While this method allows to visually compare several histograms, the overlap score used to measure the discrepancy between two distributions is sensitive to symmetries and thus it might be an inaccurate metric. In this section we develop our own novel quality assessment measures that capture idiosyncrasies not covered by Maier et al. (2013).

First, we define a similarity coefficient based on the Pearson’s correlation coefficient. Let **A** be a *n* × *n* symmetric matrix holding the pairwise distances between all genes. In order to measure how faithfully this matrix preserves the pairwise distances with respect to another *n* × *n* distance matrix **B**, we define the following coefficient:

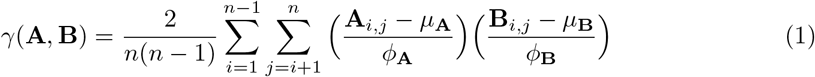

where:

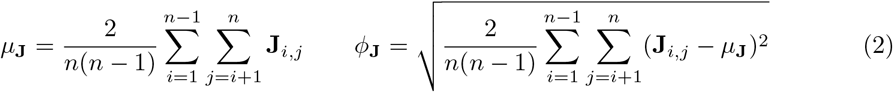

Intuitively, *γ*(**A, B**) computes the Pearson’s correlation between the elements in the upper-diagonal of matrices **A** and **B**. We leverage this similarity function in several evaluation scores.

#### 2.1.1 Distance between *real* and *artificial* distance matrices

Let **X** and **Z** be two matrices containing *m*_1_ and *m*_2_ *n*-dimensional observations that are sampled independently from the real *p*_*real*_ and synthetic *p*_*g*_ distributions, respectively. For a given distance function *d*, we define two *n* × *n* distance matrices **D**^*X*^ and **D**^*Z*^ as:

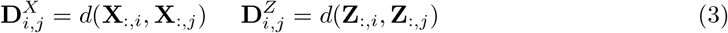

The coefficient *γ*(**D**^*X*^, **D**^*Z*^) measures whether the pairwise distances between genes from the real data are correlated with those from the synthetic data.

#### 2.1.2 Distance between *real* and *artificial* dendrograms

Let *C* : ℝ^*n*×*n*^ → ℝ^*n*×*n*^ be a function that performs agglomerative hierachical clustering according to a given linkage function, taking a *n* × *n* distance matrix as input and returning the *n* × *n* distance matrix of the resulting dendrogram. We define the *real* and *artificial* dendrogrammatic distance matrices **T**^*X*^ and **T**^*Z*^ as:

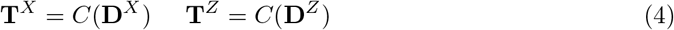

The coefficient *γ*(**T**^*X*^, **T**^*Z*^) measures the structural similarity between the dendrograms, giving a score close to 1 when the *real* and *artificial* dendrograms have a similar structure. Consequently, this metric encourages the distribution *p*_*g*_ to preserve the relationships among groups of genes that are found in *p*_*real*_.

#### 2.1.3 Squared difference between cophenetic correlation coefficients

The coefficient *γ*(**D**^*J*^, *C*(**D**^*J*^)) is known as the cophenetic correlation coefficient (Sokal and Rohlf, 1962), and it measures how faithfully a dendrogram preserves the original distance matrix. Concretely, this score quantifies the amount of information that is lost with respect to the original distance matrix when we perform hierarchical clustering on a gene expression dataset.

It is reasonable to expect the cophenetic coefficients from the real and synthetic datasets to be similar when the synthetic data is realistic. In other words, we expect the value (*γ*(**D**^*X*^,*C*(**D**^*X*^)) − *γ*(**D**^*Z*^,*C*(**D**^*Z*^)^2^ to be close to 0 when the distribution *p*_*g*_ approximates *p*_*real*_ well. Figure 1 shows the intuition behind the distance correlation coefficients.

**Figure 1:**
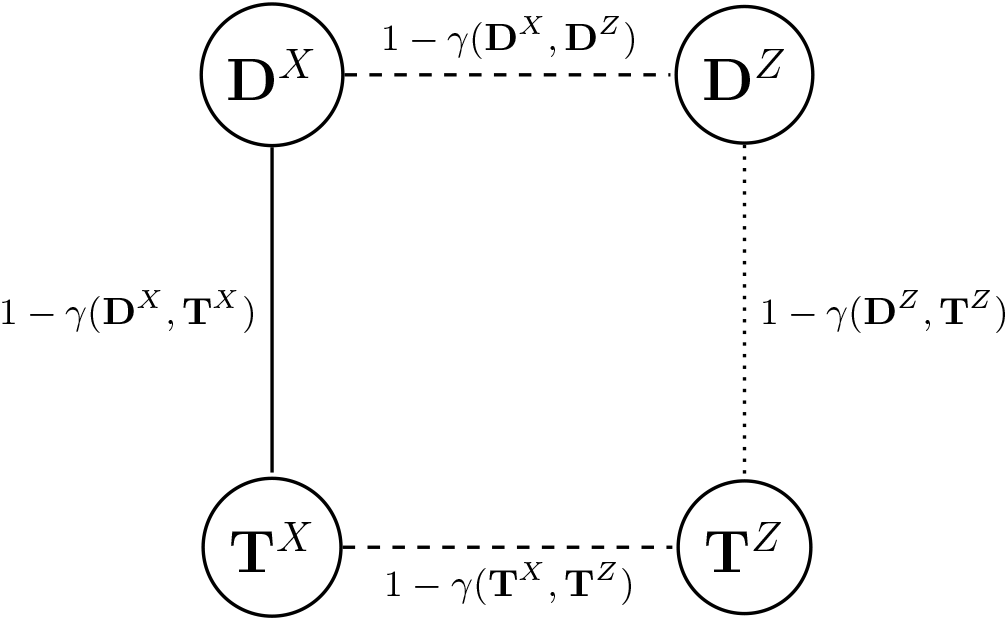
Intuition of the *γ*-based quality checks. When the synthetic data resembles the real data, the distance matrices **D**^*X*^ and **D**^*Z*^ are similar and the distance 1 – *γ*(**D**^*X*^, **D**^*Z*^) is close to 0. Likewise, 1 – *γ*(**T**^*X*^, **T**^*Z*^) is close to 0 when the synthetic and real dendrogram structures are near to each other. Even though we do not have control over 1 – *γ*(**D**^*X*^, **T**^*X*^), the distance 1 – *γ*(**D**^*X*^, **T**^*Z*^) should not deviate a lot from it when the synthetic data is realistic. Ideally, for a perfect distribution *p*_*g*_, the nodes **D**^*X*^ and **D**^*Z*^; **T**^*X*^ and **T**^*Z*^ would overlap.

#### 2.1.4 Weighted sum of TF-TG similarity coefficients

Let 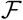 be the set of transcription factor (TF) indices, 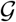 a function returning the set of indices of the target genes (TGs) that are regulated by a given TF, and *w*_*f*_ a positive coefficient associated to the importance of TF *f*. We define the following coefficient:

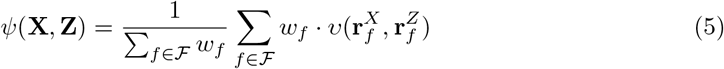

where the vector 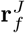 is constructed as:

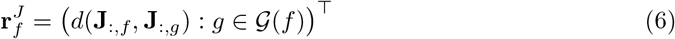

and *v*(**a, b**) is the cosine similarity between vectors **a** and **b**:

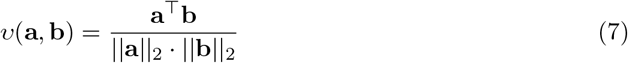

The coefficient *ψ*(**X, Z**) measures whether the TF-TG dependencies in the synthetic data resemble the ones from the real data. It is bounded as −1 ≤ *ψ*(**X, Z**) ≤ 1, and a value close to 1 indicates that the subsets of TGs for different TFs have the same general direction in terms of the vectors defined by their gene expression values. The coefficients *w*_*f*_ are arbitrary, but we choose 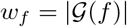 (the importance of a TF is proportional to the number of genes that it regulates).

In this case, the cosine similarity is preferred over the Pearson’s correlation coefficient *ρ* because for certain TFs (i.e. the ones regulating a small amount of genes) the vector 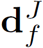 might potentially have a standard deviation 0, and as a result *ρ* would be undefined. The cosine similarity is bounded as −1 ≤ *v*(**a, b**) ≤ 1, and a value of 1 indicates that **a** and **b** point towards the same direction.

#### 2.1.5 Weighted sum of TG-TG similarity coefficients

Similarly, we define a coefficient *ϕ* to measure whether the expression of TGs regulated by the same TF in synthetic data conforms well with the analog expressions in real data:

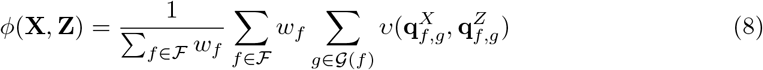

where 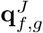 is the vector of distances between gene *g* and all the genes regulated by *f* (excluding *g*):

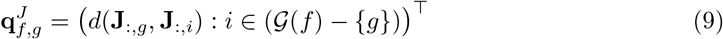

### 2.2 Adversarial simulator

We design an adversarial simulator of expression data based on a Wasserstein GAN with gradient penalty (WGAN-GP; Gulrajani et al., 2017) to design a generative model of gene expression data. In this framework, the generator *G* and the critic *D* are trained to minimize (Gulrajani et al., 2017):

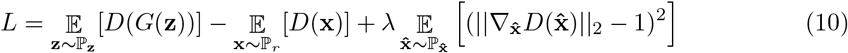

where ℙ_*r*_ is the data distribution, ℙ_**z**_ the noise distribution of the generator’s input, λ an hyperparameter, and 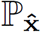 a uniform distribution over points along straight lines that connect pairs of real and generated samples. The last term of equation 10 is a gradient penalty and enforces a 1-Lipschtiz constraint on the critic. To produce synthetic gene expression examples, we first train the adversarial simulator using the default WGAN-GP algorithm and hyperparameters. Then, we sample **z** ~ ℙ_**z**_ and compute *G*(**z**) to generate a synthetic sample.

We use dropout (Srivastava et al., 2014) in both the critic and the generator’s networks to make them less reliant on individual neurons. In addition, we employ batch normalization (BN; Ioffe and Szegedy, 2015) in the generator to reduce internal covariate shift. However, we omit BN in the critic, as the norm of its gradients is penalized based on individual examples as opposed to entire batches (Gulrajani et al., 2017).

### 2.3 Data standardization and recovery

We standardize the expression values to keep them within a reasonable range and to make the learning process easier. Let **X** be an *m* × *n* matrix of expression values, where *m* is the number of samples and *n* is the number of genes. We standardize the data as follows:

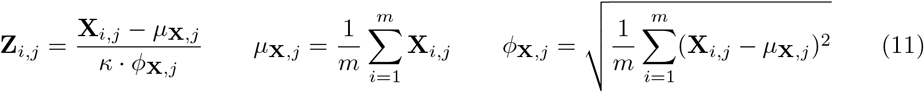

where *κ* is a user-definable constant. The expression values of each gene in the resulting standardized expression matrix **Z** have mean 0 and standard deviation *κ*^−1^.

Now, let 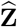 be an *m* × *n* matrix produced by a given generative model. To recover the original gene expression ranges, we first standardize the produced expression values to make them have mean 0 and standard deviation 1 for each gene:

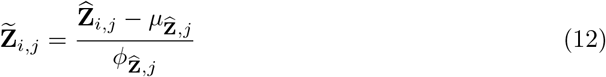

Finally, we apply the following transformation to recover the original ranges:

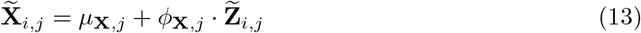

Each gene *j* has now mean *μ*_**X**,*j*_ and standard deviation *ϕ*_**X**,*j*_.

## 3 Results

### 3.1 Diversity of the generated data

Generative adversarial networks often prefer to generate data from very few modes, resulting in what is known as mode collapse (Goodfellow, 2017). Therefore, it is sensible to examine whether the diversity of gene expression data produced by gGAN is broad enough. Unlike in other domains such as image generation, where one can visually evaluate image similarity, determining whether two expression samples are similar is complicated. Here, we provide several qualitative measures that allow to verify that mode collapse does not occur.

To assess the degree of sample similarity in the synthetic dataset, we compute the Pearson’s correlation coefficient between all pairs of expression samples. Then, we compare the resulting distribution with that of real data (figure C.1). Despite being slightly shifted, the distribution for gGAN (mean sample correlation=0.74, sd= 0.14) is reasonably close to the target distribution (mean sample correlation=0.72, sd= 0.13), suggesting that gGAN produces samples coming from a reasonable amount of modes. To further show that the arrays are visually different from each other, in figure 2 we provide an overview of the real and synthetic datasets, where we perform hierarchical clustering both at sample and gene level.

**Figure 2:**
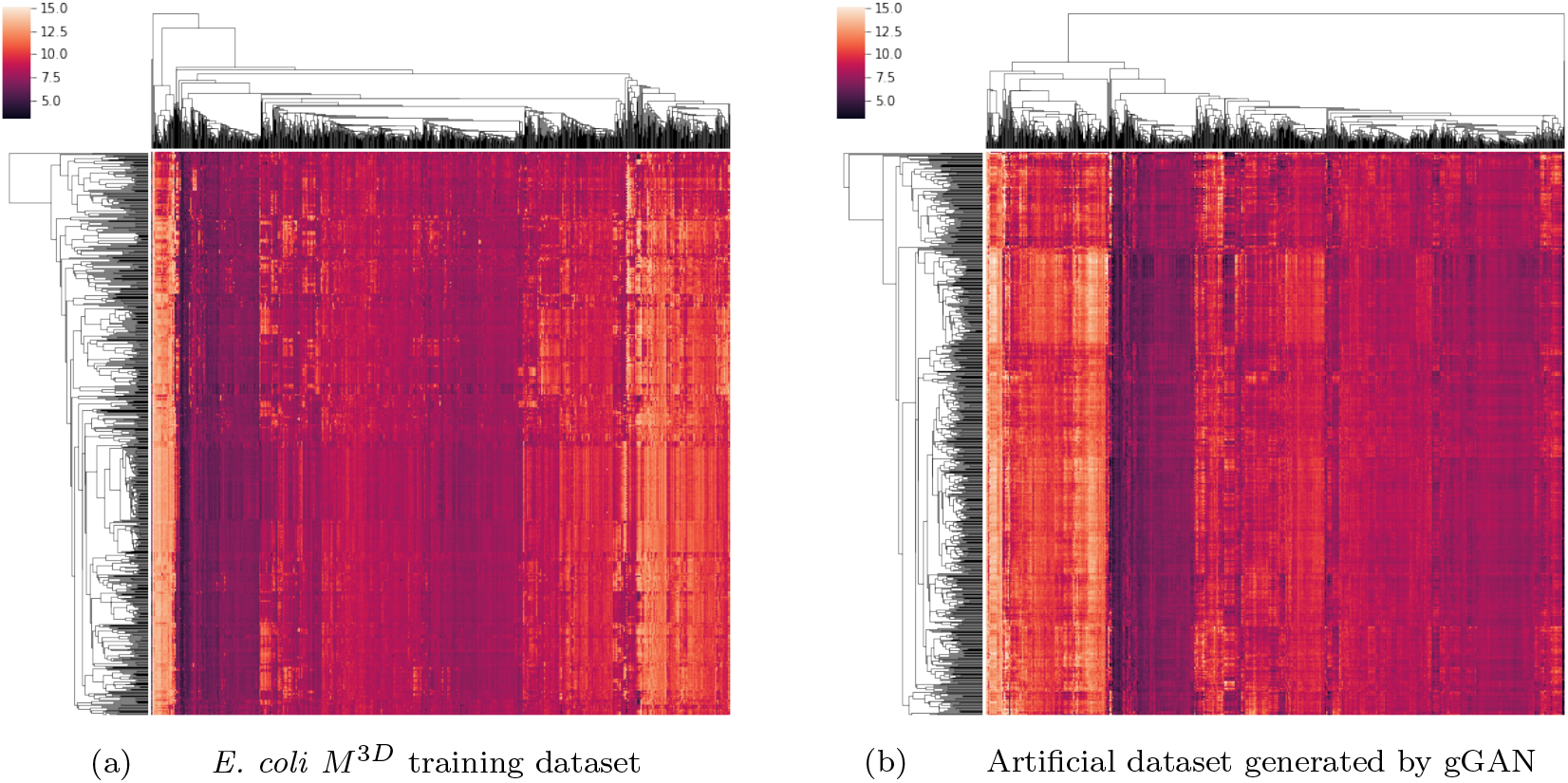
Clustering gene expression data from the *E. coli M*^3*D*^ and the gGAN datasets (rows: samples, columns: genes) for the CRP hierarchy of genes (1076 genes; see section A.2.1). We rearrange the expression matrix to enforce related genes to be adjacent to each other. Samples are also reordered according to the resulting dendrogram of samples.

Additionally, we analyze how well the individual gene distributions are preserved. In figure 3 we show the synthetic distributions of individual genes, including master regulators in the first row, and we compare those with the real gene distributions. For each gene, we also show the correlation between the real and synthetic inter-gene distance vectors and we compare them with an approximate upper bound given by the real data. Our results suggest that gGAN is able to capture the different states and modes in which these genes express and that the gene interactions are reasonably well-preserved.

**Figure 3:**
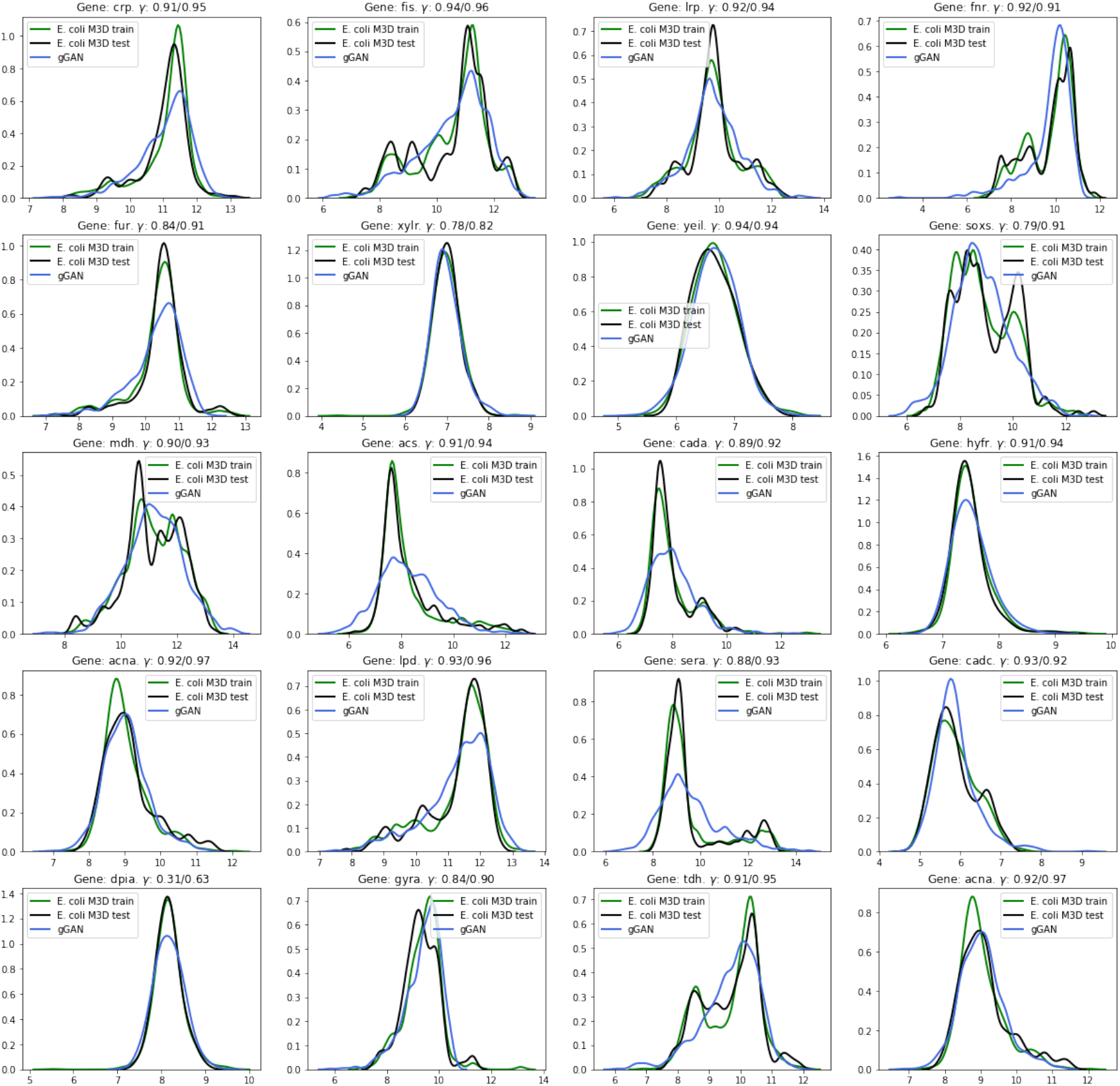
Overview of the gene distributions from a dataset (680 samples) produced by gGAN (x-axis: log expression. y-axis: density). The first row corresponds to the univariate distributions of master regulator genes. Genes in the next rows are directly regulated by each master regulator in the given column (among others), and are chosen randomly among the 4297 genes. Let **D**^*X*^ and **D**^*Z*^ be two 4297 × 4297 matrices corresponding to the gene distance matrices (as defined in section 2.1.1) of the *E. coli M*^3*D*^ test set and the gGAN dataset, respectively. Then, coefficient *γ* measures the Pearson’s correlation between 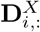 and 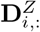 for any given gene *i*. The second value indicates the approximate upper bound for *γ*, obtained when **D**^*Z*^ is set to be the distance matrix from the *E. coli M* ^3*D*^ train set.

### 3.2 Comparison with existing methods

We compare our approaches with other existing methods: SynTReN (Van den Bulcke et al., 2006) and GeneNetWeaver (Schaffter et al., 2011). Given a GRN, these two methods model gene regulatory interactions with ordinary and stochastic differential equations based on Michaelis-Menten and Hill kinetics. SynTReN does not support GRNs with loops. To provide a fairer comparison, we opt for testing the performance on the subset of genes from the cAMP receptor protein (CRP) hierarchy, prioritizing outgoing edges from genes in top levels of the hierarchy over edges coming from bottom levels (see algorithm 1).

We generate a gene expression dataset of 680 samples both for SynTReN and GNW. For both methods, we create a network with 1076 nodes (without background nodes) and we connect them according to the forementioned CRP hierarchy. For SynTReN, we use the default parameters (probability for complex 2-regulator interactions: 0.3; biological noise: 0.1 out of 1; experimental noise: 0.1 out of 1; noise on correlated inputs: 0.1 out of 1) and CRP as the only external node. For GNW, we choose to model gene interactions with stochastic differential equations (coefficient of noise term: 0.05), and we produce multifactorial experiments using the default settings for the DREAM4 network inference challenge^1^.

#### 3.2.1 Analysis of synthetic data

Here we provide an analysis on the quality of the data generated by SynTReN, GNW and gGAN for the CRP hierarchy. To establish approximate lower and upper bounds to our evaluation metrics, we also analyze the results for a random simulator and an idyllic simulator based on the real samples from the *E. coli M*^3*D*^ train set, respectively.

SynTReN and GNW produce normalized log expression values ranging from 0 to 1 and neither SynTReN nor GNW specify how to rescale the resulting data. Maier et al. (2013) suggest multiplying the artificial ranges by a factor to make the median of the real and artificial gene range histograms match. However, in the GNW analysis by Maier et al. (2013), the absolute gene expression levels of *E. coli* and their gene ranges do not match. We attribute this to the fact that the GNW data that they generate is not properly rescaled a posteriori. Here, in contrast, we show that the rescaling procedure described in section 2.3 allows to accurately match the overall gene expression values and ranges for all three methods (figures 4 and 5). It is also worth noting that this procedure does not significantly compromise the performance of SynTReN and GNW on the upcoming evaluation checks, as these analyses are all based on correlation coefficients and thus they are not sensitive to the gene scales.

**Figure 4:**
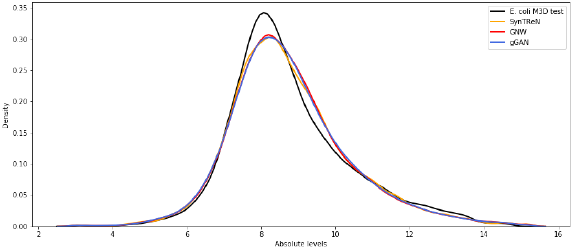
Distribution of gene intensities.

**Figure 5:**
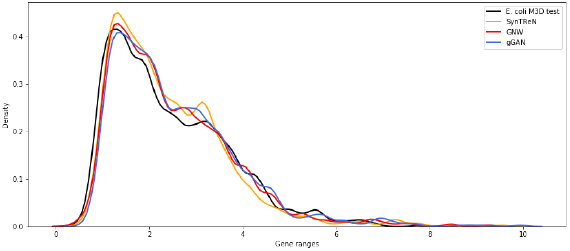
Range of gene expressions.

The background distributions of the Pearson’s correlation coefficients are considerably different. On the one hand, the distribution for SynTReN is bimodal and genes are either highly negatively correlated or highly positively correlated among each other. On the other hand, the distribution for GNW is significantly more peaked than the real distribution. This could be due to an exaggerated amount of noise being added to the generated data, which reduces the absolute correlation among pairs of genes. In contrast, the background distribution for gGAN is significantly closer to the target distribution, but there there are opportunities for further enhancements. These results notoriously compromise the TF-TG and TG-TG histograms (figures C.2 and C.3), from which we can not draw any meaningful conclusions.

IN figure 7 we illustrate the histograms of the activity of TFs. They are formed by computing the fraction of samples in which TF targets exhibit rank differences with respect to other non TF targets, according to a two-sided Mann-Whitney rank test. These tests are corrected with the Benjamini-Hochberg’s procedure in order to account for multiple testing and reduce the false discovery rate. We observe that, for all three simulators, a large amount of TFs are found inactive in every chip. However, for GNW and SynTReN the density of TFs that are active in a significant portion of chips is considerably smaller than for the *E. coli M* ^3*D*^ test set. Although the TF activity histogram is specific to the *M*^3*D*^ dataset (and not necessarily to the real *E. coli* distribution), we find these densities rather small. Conversely, the overall TF activity of *M*^3*D*^ in these regions is better preserved in the dataset generated by gGAN.

**Figure 6:**
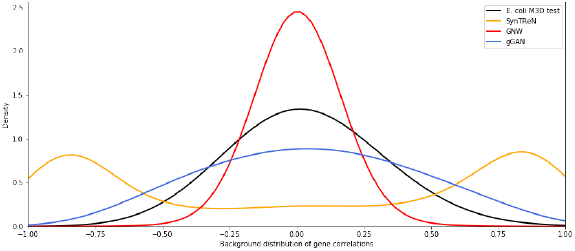
Background distribution of the correlation coefficients between all pairs of genes.

**Figure 7:**
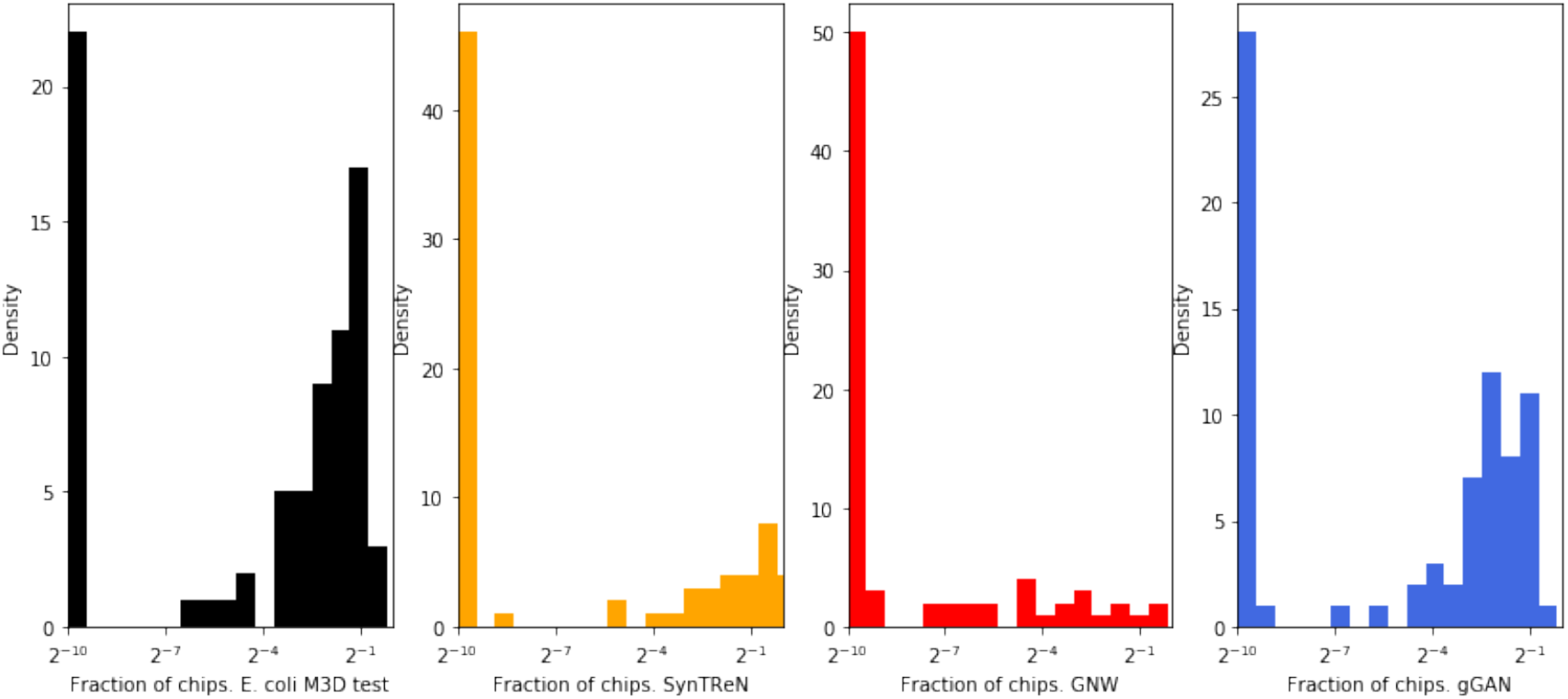
Histograms of the TF activity.

Table 1 shows a quantitative comparison of the three methods, including the lower and upper bounds given by the random and real simulators, respectively. We observe that gGAN closely approximates the upper bound in every metric, outperforming SynTReN and GNW by a large margin. In fact, SynTReN and GNW are not much better than the random simulator in terms of the realism of the generated data. We attribute this mainly to the fact that SynTReN and GNW rely exclusively on the source GRNs to produce synthetic data. In addition, this shows that the linear system of ODEs/SDEs defined via Michaelis-Menten and Hill equations are not enough for modelling gene dependencies in spite of being theoretically well-grounded. In contrast, gGAN leverages real expression data to build a generative model in an unsupervised manner, but does not require any information on the regulatory interactions. By playing a two-player game, the critic acts as a teacher that instructs the generator on how to optimize its parameters in order to produce realistic gene expression data, wherein the way in which genes interact with each other is preserved. Furthermore, our results show that the GAN trained on the full set of genes (gGAN_1_; section B) is also capable of generating realistic expression data for a subnetwork of genes.

**Table 1:**
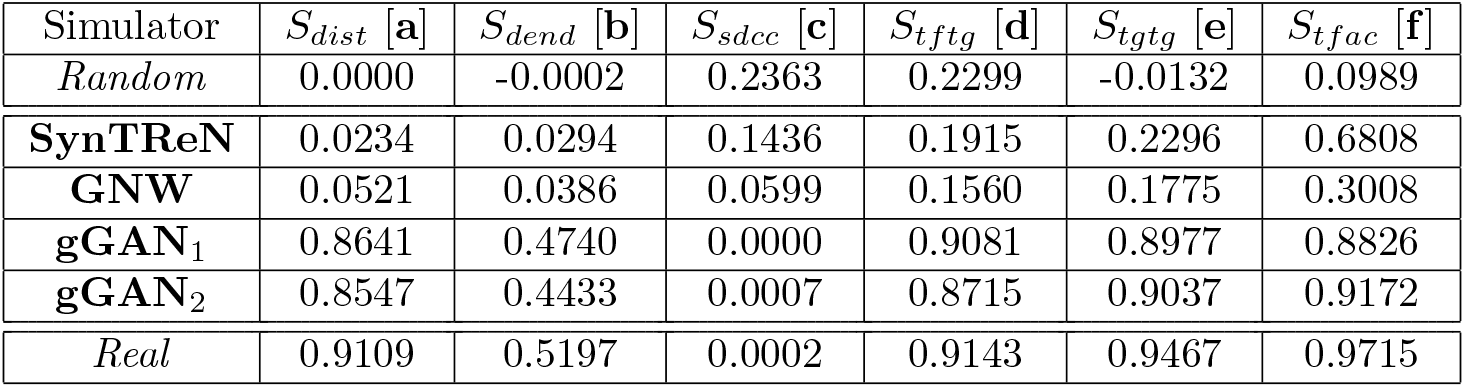
Quantitative assessment of the generated data. We include the results for a *random* and a *real* (*M*^3*D*^ train) simulators. Let **X** be the test set, and **Z** the matrix of observations sampled from any given simulator. **gGAN**_1_ is trained on all genes (section B). **gGAN**_2_ is trained on the CRP hierarchy. [**a**] *S*_*dist*_ = *γ*(**D**^*X*^,**D**^*Z*^). [**b**] *S*_*dend*_ = *γ*(**T**^*X*^,**T**^*Z*^). [**c**] *S*_*sdcc*_ = (*γ*(**D**^*X*^,**T**^*X*^) - *γ*(**D**^*Z*^,**T**^*Z*^))^2^. [**d**] *S*_*tftg*_ = *ψ*(**D**^*X*^,**D**^*Z*^). [**e**] *S*_*tgtg*_ = *ϕ*(**D**^*X*^,**D**^*Z*^). [**f**] *S*_*tfac*_ = *ω*(**X, Z**). For the three latter measures, the importance of each TF is proportional to its number of TGs.

## 4 Conclusion

Here we have studied the problem of generating realistic *E. coli* gene expression data. We have divided this problem into two main tasks: assessing the realism of a dataset and building a generative model to produce realistic gene expression data.

For the first task, we have developed our own novel evaluation scores. These metrics allow to accurately quantify the discrepancies of several statistical properties between the synthetic and real data distributions. Moreover, to provide a qualitative analysis of the simulated expression data, the histograms proposed by Maier et al. (2013) have allowed us to visualize and interpret several meaningful properties.

As a result of our analysis, we have shown that existing simulators fail to emulate key properties of gene expression data, as pointed out by Maier et al. (2013). In particular, one of the most surprising results is that SynTReN and GNW poorly preserve the properties derived from gene regulatory networks such as the TF-TG interactions. This is undesirable, as these simulators are specifically designed for the purpose of benchmarking network inference algorithms, and the goal of the linear systems of ODEs/SDEs is precisely to simulate relationships between TFs and TGs.

For the second task, we have implemented a simulator based on a Wasserstein Generative Adversarial Network with gradient penalty (Gulrajani et al., 2017). To our best knowledge, GANs have not previously been applied to build a realistic simulator of gene expression data. To enable learning from a high-dimensional dataset with a scarce number of samples and keep the complexity of our model under control, our training algorithm incorporates several regularization mechanisms such as dropout and batch normalization.

After carefully optimizing our adversarial model, the resulting simulator generates expression data with a high degree of realism according to our evaluation metrics. We have shown that our approach outperforms existing simulators by a large margin in terms of the realism of the generated data, and the quality scores of our simulator are reasonably close to the upper bound given by the real data. More importantly, our results show that gGAN is, to our best knowledge, the first gene expression simulator that passes the Turing test for gene expression data employed by Maier et al. (2013).

## Software

The code is available at: https://github.com/rvinas/adversarial-gene-expression

## Footnotes

1 http://gnw.sourceforge.net/dreamchallenge.html

