## Supplementary material for "Adversarial generation of gene expression data"

### Appendix

### A Materials

In this study, we leverage data from *Escherichia coli* (*E. coli*) to design a generative model of its expression data. This bacterium has been especially useful to molecular biologists because of the ease with which it can be studied in the laboratory (Cooper, 2000). In addition, *E. coli* has a relatively small and simple genome (∼4,400 genes) and its gene expression mechanisms are relatively well understood (Salgado et al., 2006).

#### A.1 *E. coli* gene expression $M^{3D}$ data

Many Microbe Microarrays Database ($M^{3D}$; Faith et al., 2008) is a database that contains gene expression data for microbes such as *Escherichia coli*, *Saccharomyces cerevisiae* and *Shewanella oneidensis*. The data from $M^{3D}$ comes from single-channel Affymetrix microarray experiments that were carried out by different laboratories (Faith et al., 2008). The data was uniformly normalized using log-scale robust multi-array average (RMA; Irizarry, 2003) to reduce batch effects and make the samples comparable across conditions.

The *E. coli* $M^{3D}$ database consists of 907 chips. There are 466 unique conditions, hence some of these chips measure expressions from replicate experiments. The database provides both the 907 original measurements and the 466 expressions averaged across replicate experiments. In this study we use the data from the 907 original measurements, and we leverage the information about replicate experiments to sensibly split the data into a train and a test set. This train-test partition strategy is further discussed in section A.2.2.

Each of the 907 chips consists of 7459 probes. These probes include the expression values of genes, intergenic regions and control probes. In this study we are specifically interested in gene expression values, so we exclude intergenic regions and control probes. The resulting dataset consists of 907 samples and 4297 features corresponding to the expression values of *E. coli* genes. The log-scaled RMA-normalized gene expression values range from 2.94 to 15.19, with an overall mean value of 8.77.

#### A.2 *E. coli* gene regulatory interactions and RegulonDB

The gene regulatory network of *E. coli* is one of the most well-characterized transcriptional networks of a single cell. RegulonDB (Gama-Castro et al., 2016) is a database that integrates biological knowledge about the transcriptional regulatory mechanisms of *E. coli*. The database gathers information from multiple biological studies to reconstruct the structure of the *E. coli* GRN. For each regulatory interaction, RegulonDB provides the name of the TF; the gene regulated by the TF; the regulatory effect (activation and/or repression); the experimental evidences that support the existence of the regulatory interaction; and a measure of the evidence strength (weak, strong or confirmed).

This study leverages information from RegulonDB for two purposes. The first one is motivated by the fact that the number of samples (907) of the *E. coli* $M^{3D}$ dataset is rather scarce in comparison to the number of genes (4297). In this work, as one of our experiments, we use RegulonDB to select a meaningful subset of *E. coli* genes by choosing a hierarchy of genes whose transcription is directly or indirectly regulated by one of the *E. coli* master regulators. The details about our selection of genes are provided in section A.2.1. Secondly, we use RegulonDB to characterize several statistical properties of the simulated datasets, with the aim of comparing them with those of the real dataset.

##### A.2.1 Selecting the CRP hierarchy

As one of the experiments in our study, we select a meaningful subset of *E. coli* genes whose expression is directly or indirectly regulated by the master regulator cAMP receptor protein (CRP). CRP regulates global patterns of transcription in response to carbon availability, and it is one of the best characterized global transcriptional regulators in *E. coli.* This receptor increases the levels of promoter occupancy by binding to the DNA sites in the promoters of its target genes, and by interacting with RNA polymerase to activate their transcription (Grainger and Busby, 2008).

Information from the *E. coli* gene regulatory network is used to extract the regulatory hierarchy in which CRP is the root node. Concretely, we use algorithm 1 with the RegulonDB network of regulatory interactions to select the CRP hierarchy from the *E. coli* $M^{3D}$ dataset. Note that, when we break loops, the resulting structure is a directed acyclic graph, as edges going from children to ancestors are ignored.

**Algorithm 1:** Selecting a hierarchy of genes. Returns the sets of nodes and edges in the hierarchy.

**Data:** We are given a set of genes $\mathcal{G}$, a root gene $g_r \in \mathcal{G}$, a network $\mathcal{S}$ of gene regulatory interactions, and a boolean $b$ indicating whether to break loops.

Initialize sets of edges, previous and current included nodes, and ancestors of each node:
    $\mathcal{E} \leftarrow \{\}$
    $\mathcal{V}' \leftarrow \{\}$
    $\mathcal{V} \leftarrow \{g_r\}$
    $\mathcal{A}(g) \leftarrow \{g\} \ \ \forall g \in \mathcal{G}$
**while** $\mathcal{V} \neq \mathcal{V}'$ **do**
    Update previous nodes:
        $\mathcal{V}' \leftarrow \mathcal{V}$
    Select nodes to be added:
        $\mathcal{C} \leftarrow \{g_1 \in \mathcal{G} \mid \exists g_2 \in \mathcal{V}, (g_2 \to g_1) \in \mathcal{S}, \neg b \vee \neg g_1 \in \mathcal{A}(g_2)\}$
    Add them to set of nodes:
        $\mathcal{V} \leftarrow \mathcal{V} \cup \mathcal{C}$
    **for** *each* $g_1 \in \mathcal{C}$ **do**
        Find parents of $g_1$:
            $\mathcal{P} \leftarrow \{g_2 \in \mathcal{V}' \mid (g_2 \to g_1) \in \mathcal{S}, \neg b \vee \neg g_1 \in \mathcal{A}(g_2)\}$
        Add edges coming from parents:
            $\mathcal{E} \leftarrow \mathcal{E} \cup \{g_2 \to g_1 \mid g_2 \in \mathcal{P}\}$
        Update ancestors:
            $\mathcal{A}(g_1) \leftarrow \mathcal{A}(g_1) \cup \mathcal{P} \cup \bigcup_{p \in \mathcal{P}} \mathcal{A}(p)$
**return** $\mathcal{V}$, $\mathcal{E}$

##### A.2.2 Train-test split

Unlike in other domains where the model's target or the evaluation metrics are clearly defined for each sample, the need for a test set in this unsupervised problem is questionable. One might argue that reporting the generalization results on a separate test set is not necessary, as we are trying to approximate the real data distribution and the train/test data are both sampled from this distribution.

However, we opt for having a test set for two reasons. First, the number of samples of the *E. coli* $M^{3D}$ dataset is rather scarce, and thus they might not represent the real distribution of *E. coli* gene expressions well enough. Therefore, comparing the properties of our generated data against those of an unseen test set might provide us with a more realistic view of our generalization capability. Second, keeping a separate test set allows to compare its properties with those of the training set and this gives an upper bound for our evaluation metrics.

We split the *E. coli* $M^{3D}$ data into train (680 samples) and test (227 samples) sets. We partition the data in a random manner, constraining replicate samples to lie within the same set. This prevents the leakage of information between sets. We use the train set to train our generative model and tune its hyperparameters, whereas the test set is kept apart and used only to report the final results.

### B Evaluating gGAN gene expression data

Here we leverage Maier's histograms (Maier et al., 2013) and our own novel quality scores to evaluate the realism of the synthetic data generated by gGAN for the full set of *E. coli* $M^{3D}$ (4297 genes). We examine the properties using the test samples as a representation of the true data distribution. Additionally, we contrast the forementioned properties between the training and the test sets. Since both sets are sampled from the real distribution, this comparison should provide us with an overview of the irreducible error and an approximate upper bound on our evaluation metrics.

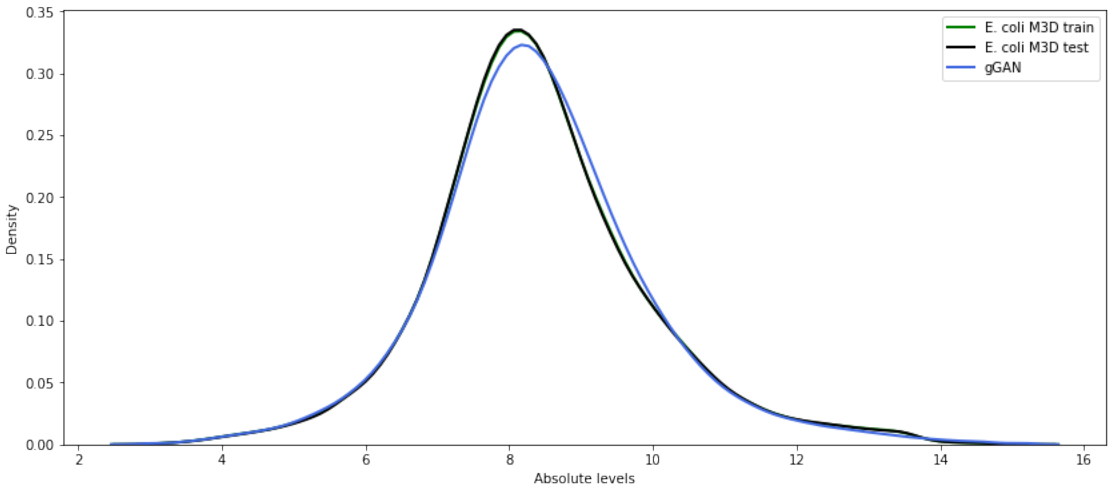


Figure B.1: Distribution of gene intensities.

Table 2: Quantitative assessment of the generated data. Let $\mathbf{X}$ be the test set, and $\mathbf{Z}$ the matrix of observations sampled from any given simulator. [**a**] $S_{dist} = \gamma(\mathbf{D}^X, \mathbf{D}^Z)$. [**b**] $S_{dend} = \gamma(\mathbf{T}^X, \mathbf{T}^Z)$. [**c**] $S_{sdcc} = (\gamma(\mathbf{D}^X, \mathbf{T}^X) - \gamma(\mathbf{D}^Z, \mathbf{T}^Z))^2$. [**d**] $S_{tftg} = \psi(\mathbf{D}^X, \mathbf{D}^Z)$. [**e**] $S_{tgtg} = \phi(\mathbf{D}^X, \mathbf{D}^Z)$. To form the dendrograms, we use agglomerative hierarchical clustering with Pearson distance and complete linkage. For the three latter scores, the importance of each TF is set to be proportional to its number of TGs.

| Simulator | $S_{dist}$ [**a**] | $S_{dend}$ [**b**] | $S_{sdcc}$ [**c**] | $S_{tftg}$ [**d**] | $S_{tgtg}$ [**e**] |
|---|---|---|---|---|---|
| *Random* | 0.0002 | 0.0003 | 0.2426 | 0.2711 | -0.0026 |
| **gGAN** | 0.8925 | 0.3344 | 0.0009 | 0.9084 | 0.9065 |
| *Real* | 0.9252 | 0.3565 | 0.0008 | 0.9228 | 0.9515 |

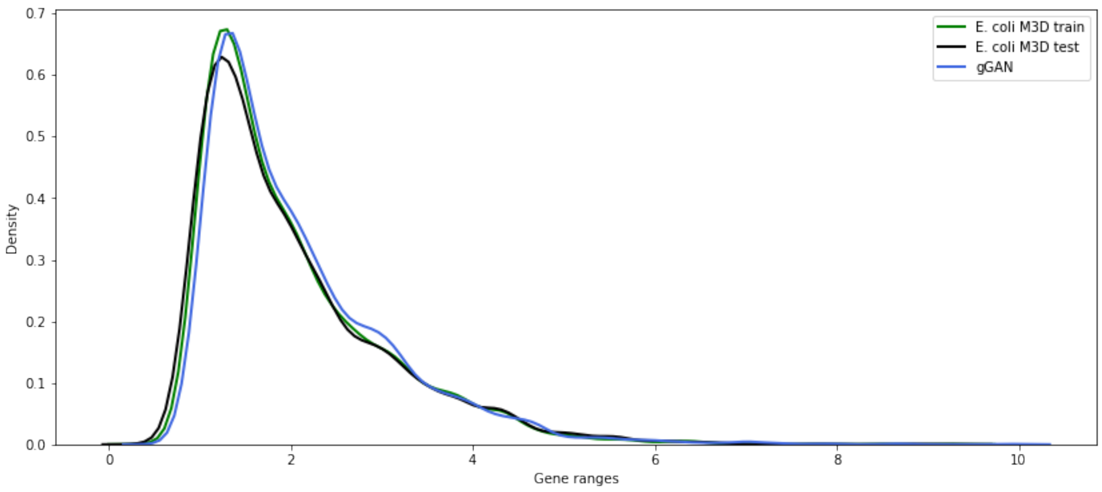


Figure B.2: Range of gene expressions.

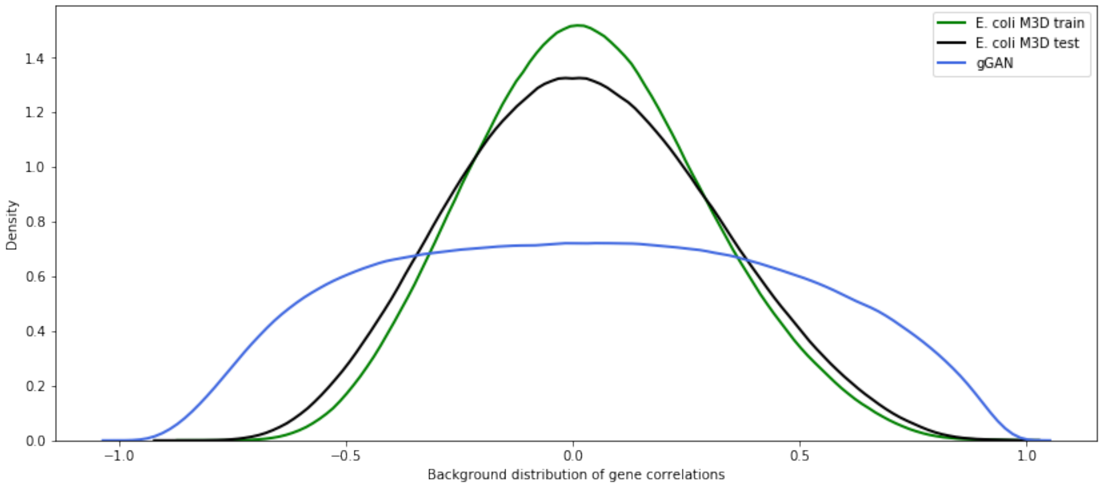


Figure B.3: Background distribution of correlation coefficients between all pairs of genes.

The gene intensities and gene ranges distributions illustrated in figures B.1 and B.2 show that gGAN is able to closely match the overall gene expression values and ranges of the real *E. coli* $M^{3D}$ data. In parallel, in figure 3 we observe that the true mean and standard deviation of each gene are preserved. This is mostly due to the rescaling procedure (section 2.3) applied at the output of the generator, as the GAN is not directly trained to match the univariate gene distributions, which are specific to the $M^{3D}$ dataset, but not necessarily to the *E. coli* real distribution of gene expressions. Concretely, the generator's loss penalizes a single sample for not being realistic rather than punishing a batch of samples for not matching the $M^{3D}$ univariate distributions. In other words, the GAN is specifically optimized to preserve the way in which genes interact with each other. Additionally, the fact that the real *E. coli* univariate distributions are generally more peaked than those of gGAN explains the small discrepancy between the overall gene intensities around the mean.

The background distribution depicted in figure B.3 reflects the overall histogram of correlation coefficients among genes. This distribution shows that the majority of genes are uncor-

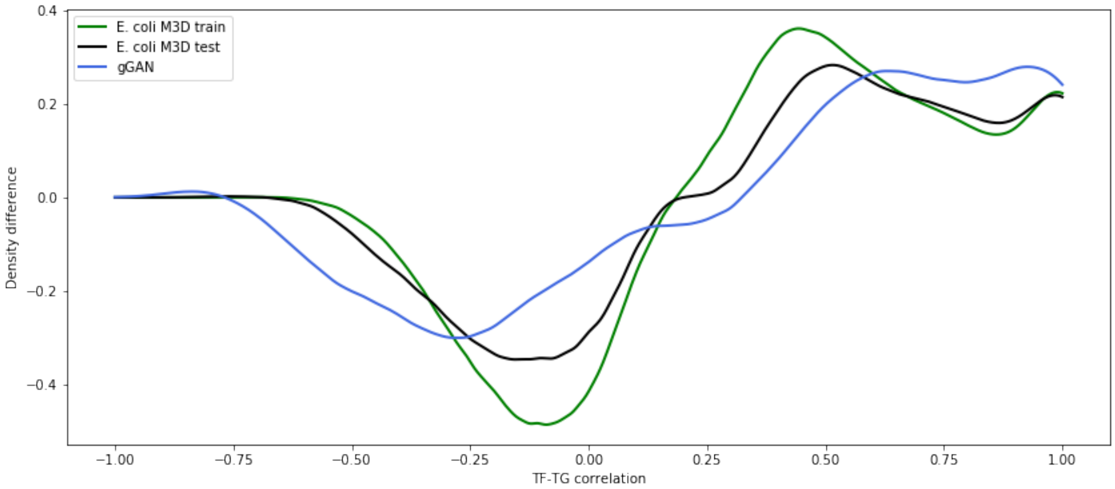


Figure B.4: Histogram of TF-TG interactions. It shows to what extent TF-TG pairs are enriched (> 0) or depleted (< 0) with respect to the background distribution.

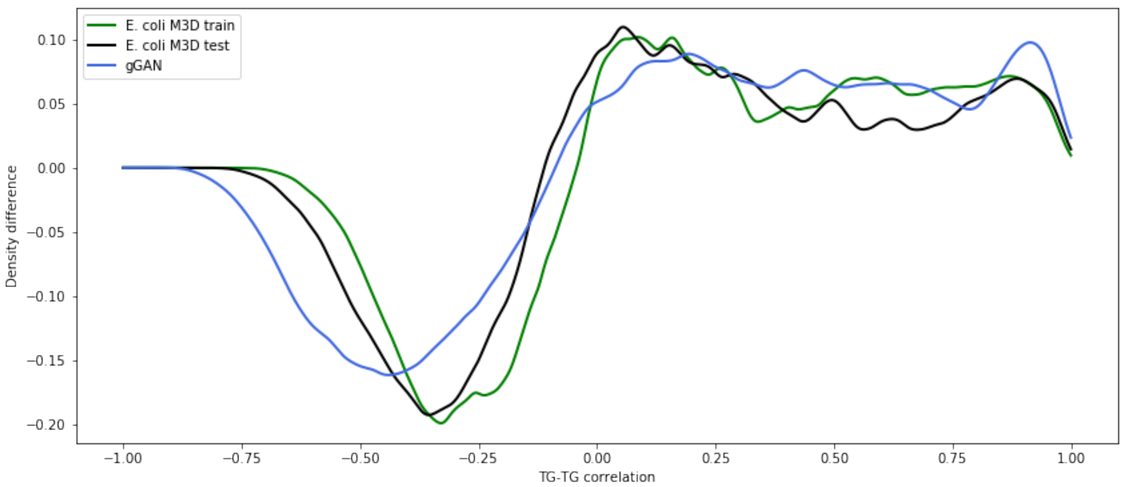


Figure B.5: Histogram of TG-TG interactions. It shows to what extent TG-TG pairs are enriched (> 0) or depleted (< 0) with respect to the background distribution.

related or slightly correlated, whereas values around the tails correspond to pairs of genes that exhibit a strong negative (left tail) or positive (right tail) correlation. Additionally, we show the same distribution on

Note that, in the background distribution, a high correlation between two genes does not necessarily imply a TF-TG regulatory interaction between them, as they could both be regulated by a common ancestor. Instead, we summarize to what extent TF-TG pairs are enriched or depleted in figure B.4. A positive or a negative difference indicates that the probability of detecting a true TF-TG interaction in the given interval is increased or decreased, respectively, with respect to the background distribution. Similarly, figure B.5 summarizes the same property for pairs of TGs regulated by the same TF. In both cases, gGAN is able to match these density differences (using the synthetic background distribution as reference) reasonably well.

In figure B.6 we examine whether the overall activity of TFs in the samples generated

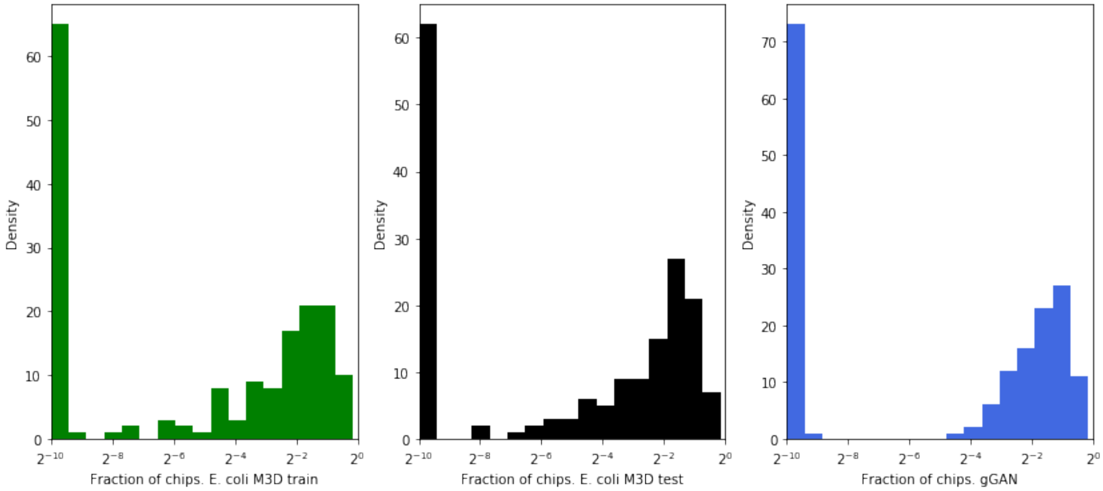


Figure B.6: Histograms of the TF activity.

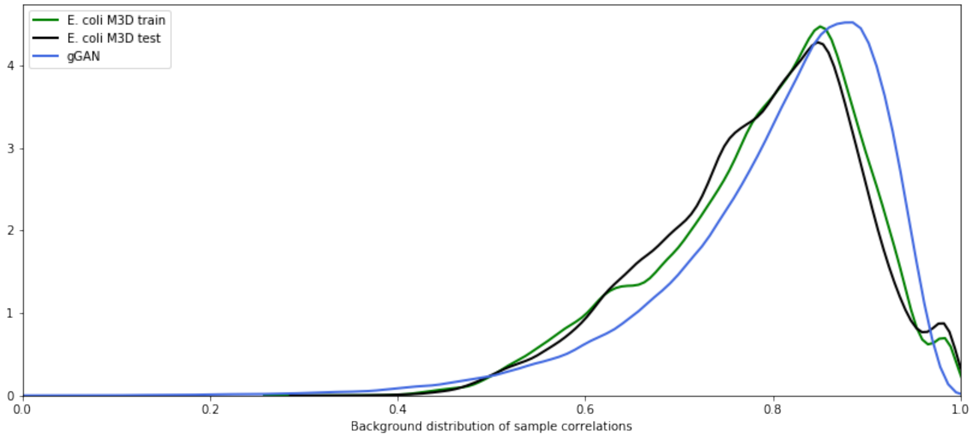


Figure B.7: Background distribution of correlation coefficients between all pairs of samples.

by gGAN resembles the levels of TF activity in the $M^{3D}$ data (Maier et al., 2013). Since the activity of a TF depends on the circumstances under which each sample was measured, this property is specific to the $M^{3D}$ dataset and thus it does not necessarily represent the true distribution $p_{data}$. Nonetheless, we observe a common trend regarding the TF activity between the $M^{3D}$ and the gGAN datasets: a large amount of TFs are found inactive in every chip, whereas a considerable amount of TFs are active in a significant portion of chips.

Finally, our most meaningful results are shown in table 2. These measures provide a quantitative summary on several gene expression properties along with an approximate upper bound on them. First, $S_{dist}$ indicates to which extent the pairwise gene distances are preserved in the data generated by gGAN. In figure 3 we show a finer-grained version of this coefficient for each gene. $S_{dend}$ measures whether the gene clusters formed in the gGAN data are similar to those from the real data. $S_{sdcc}$ shows the squared difference between the cophenetic coefficients of the *E. coli* $M^{3D}$ and gGAN expression matrices (lower is better). Lastly, $S_{tftg}$, $S_{tgtg}$ and $S_{tfac}$ summarize whether the RegulonDB (Gama-Castro et al., 2016) *E. coli* TF-TG, TG-TG and TF activity correlations are preserved in the gGAN dataset, respectively.

### C Supplementary figures

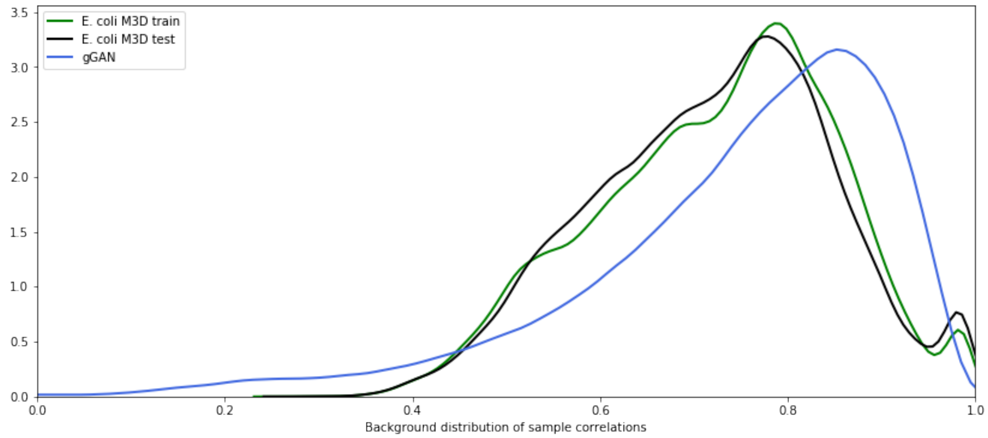


Figure C.1: Background distribution of the Pearson's correlation coefficients between all pairs of samples, for the gGAN trained on the CRP hierarchy. The mean sample correlation are 0.72 (sd 0.13) and 0.74 (sd 0.17) for the real and simulated datasets, respectively.

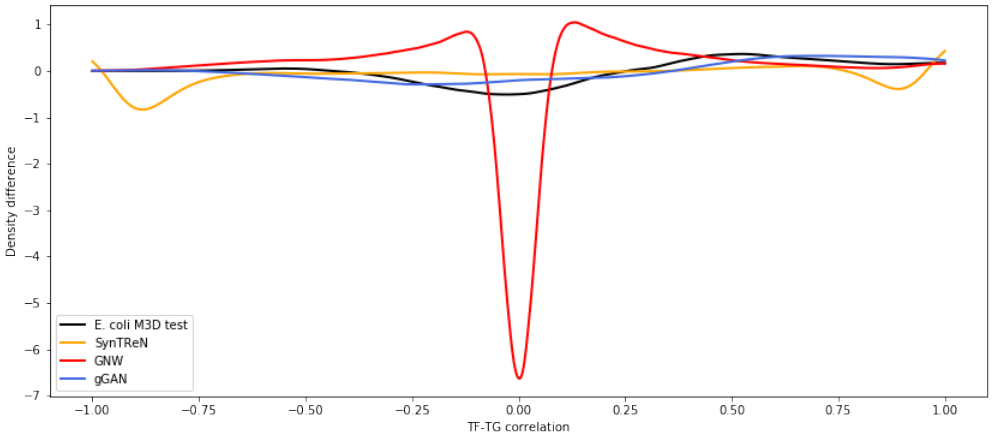


Figure C.2: Histogram of TF-TG interactions, for the gGAN trained on the CRP hierarchy. It shows to what extent highly correlated TF-TG pairs are enriched (> 0) or depleted (< 0).

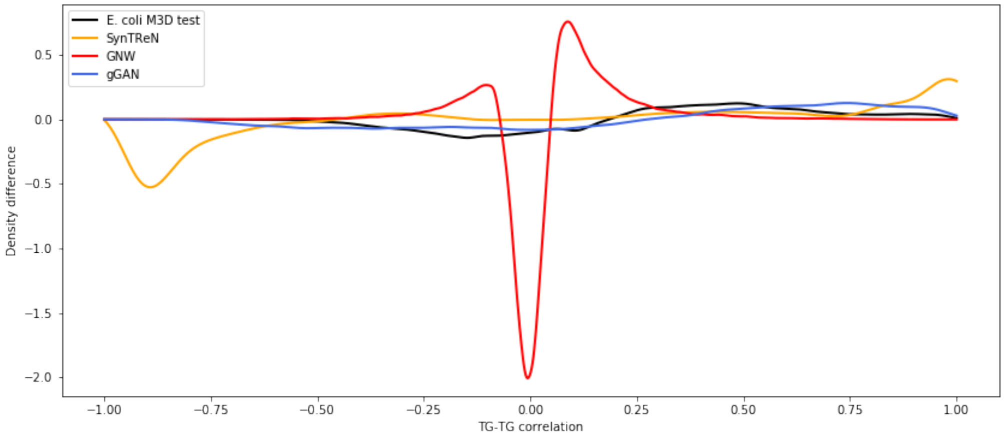


Figure C.3: Histogram of TG-TG interactions, for the gGAN trained on the CRP hierarchy. It shows to what extent highly correlated TG-TG pairs are enriched ($> 0$) or depleted ($< 0$).
